# Building subtraction operators and controllers via molecular sequestration

**DOI:** 10.1101/2023.04.24.538183

**Authors:** Christian Cuba Samaniego, Yili Qian, Katelyn Carleton, Elisa Franco

**Author notes:** Authors contributed equally (Christian Cuba Samaniego, Yili Qian, Katelyn Carleton).

## Abstract

We show how subtraction can be performed via a simple chemical reaction network that includes molecular sequestration. The network computes the difference between the production rate parameters of the two mutually sequestering species. We benefit from introducing a simple change of variables, that facilitates the derivation of an approximate solution for the differential equations modeling the chemical reaction network, under a time scale separation assumption that is valid when the sequestration rate parameter is sufficiently fast. Our main result is that we provide simple expressions confirming that temporal subtraction occurs when the inputs are constant or time varying. Through simulations, we discuss two sequestration-based architectures for feedback control in light of the subtraction operations they perform.

## I. Introduction

MOLECULAR sequestration (titration) is a prevalent mechanism in biology that consists of the stoichiometric binding of two species. Sequestration can involve many classes of molecules; examples include RNA-RNA binding, transcription factors binding to promoters, enzymes and their substrates, receptors and their ligands, or antibodies and antigens. This binding process plays a crucial role in various biological processes, including signal transduction, regulation of gene expression, immune response, and metabolic pathways. Is there an underlying operation being computed by molecular sequestration, that is key to enable a broad variety of functions? This question is difficult to answer in the native context of the cell, but there is evidence that sequestration enables the computation of the difference in the level of participating molecules [1], [2]. In synthetic biology, mathematical modeling has shown how to leverage this property to build molecular controllers [2]–[6], networks for frequency modulation [7], bistable switches with tunable hysteresis [8], [9], gradient detectors [10], [11], and biomolecular perceptrons [12]; some of these ideas have been experimentally demonstrated [13]– [18]. It is abundantly evident that sequestration helps compute the difference of two inputs *at steady-state* [1]–[4], [17], and linear analysis shows that it works as a sum junction [19]. However, a complete analysis of the dynamical behavior of these sequestration reactions is still missing, making it challenging to understand/design its behavior when the inputs are time-varying and in the presence of dilution/degradation of the reactants.

Another limitation that needs to be addressed is the fact that sequestration can only calculate the *positive part of a subtraction operation*, due to the biological nature of the participating components [20]. In this sense, it bears similarity to functions computing an absolute value [7].

Here we provide a complete temporal analysis of a molecular sequestration network, modeled through ordinary differential equations, and we show that it can compute differences in the presence of dilution/degradation, both transiently and at steady state. To come to this conclusion, we introduce a change of variables that makes it easier to apply time scale separation arguments, which decouples the system into a linear and non-linear part. This allows us to find an approximate solution of the output of this network, which is proportional to the difference of the molecular inputs, whether they are constant or time varying. We then discuss how sequestration can be used as a low-pass controller within biomolecular feedback loops. Through simulation we also show how low-pass controllers in a “dual rail” architecture improves the response speed of a feedback circuit tracking time-varying inputs. We expect that our modeling results will boost the use of sequestration within complex synthetic biological networks, and that they will facilitate the formulation of hypotheses about the roles of sequestration in natural systems.

### II. A model for molecular sequestration and its singular perturbation form

Our chemical network for subtraction includes two species, *X* and *Y*, that mutually sequester and bind to form a complex *C* (Fig. 1). We assume that *X* and *Y* have zero-order time-varying production rates *α*(*τ*) and *β*(*τ*). We model dilution/degradation as a first-order process with rate parameter *ϕ*, assumed to be the same for simplicity. The sequestration of *X* and *Y* happens with a constant rate parameter *γ*. We postulate that *γ* can be rescaled [3], [5]. The chemical reactions are:

**Fig. 1.**
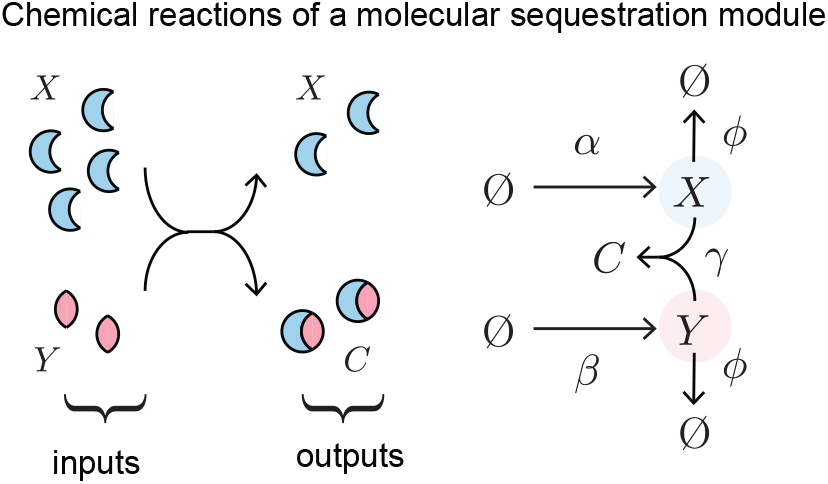
Schematic of the reactions occurring in the molecular sequestration network considered here

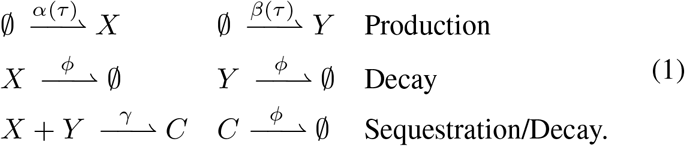

From the reactions in (1), we derive an Ordinary Differential Equation (ODE) model using the law of mass action:

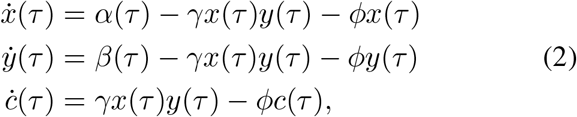

where the 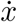, 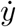, and *ċ* indicate derivatives with respect to the time variable *τ*. Previous work considering similar reactions assumed (i) *ϕ* = 0 and (ii) *α*(*τ*) *> β*(*τ*) for all time [2].

When *α* = *β* = 0 we can immediately provide intuition as to why molecular sequestration realizes a subtraction operator. First, sequestration implies there is a 1:1 binding ratio between *X* and *Y* (one molecule of *X* binds to one molecule of *Y*). Then, in the limit case where *ϕ →*0, the final concentration of the non-limiting species is the difference between the initial concentrations of the sequestering species [1]. Suppose *x*(0) *> y*(0): the final concentration of *X* (non-limiting species) will be equal to the difference in the initial concentrations *x*(0) −*y*(0), given that all free *Y* (the limiting species) is bound into the complex *C*. In practice, however, matters are made more complex due to the presence of time varying input functions *α*(*τ*) and *β*(*τ*), and of degradation and dilution, scaled by parameter *ϕ*. Next, we illustrate how to examine this case by introducing an approximation valid for constant and time-changing inputs, which we assume to be bounded and non-negative. We do not introduce requirements on the relative magnitude of the inputs.

#### Assumption 1

There exist positive constants *α*^*∗*^, *β*^*∗*^ such that 0 *≤ α*(*t*) *≤ α*^*∗*^ and 0 *≤ β*(*t*) *≤ β*^*∗*^ for all *t ≥* 0.

To obtain a non-dimensional model, we first rescale time as *t* = *ϕτ*, and define new states variables 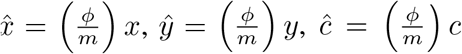, and new rates 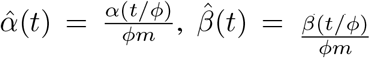, and 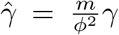, where *m* = max(*α*^*∗*^, *β*^*∗*^). Our non-dimensional model is now:

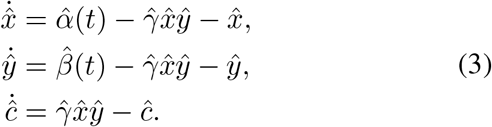

Next, we apply a coordinate transformation that makes it possible to put the system in its singular perturbation form. We begin by applying a coordinate transformation:

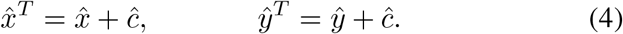

The new variables 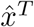 and *ŷ*^*T*^ represent the total concentration of species *X* and *Y*, which includes molecules that are free or bound to make the complex *C*, and system (3) becomes:

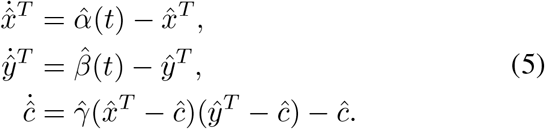

Let 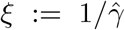, and system (5) can be put into a singular perturbation form:

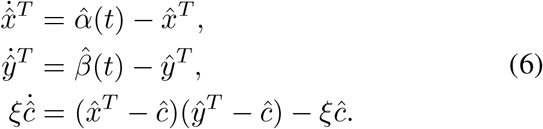

Note that the first two equations of the system above are linear and do not depend on *ξ*. In the next sections, we use the transformed system (6) to demonstrate its capacity to perform a subtraction operation.

### III. The molecular sequestration network performs a subtraction operation

We will show that the solution 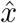 of system (3) satisfies:

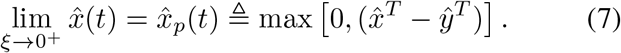

In other words, 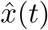 can be approximated by 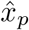, the *non-negative part of the subtraction* of variables 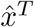 and *ŷ* ^*T*^ that capture the total amounts of molecules 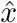 and *ŷ*. It is also possible to prove that lim _ξ →0+_ *ŷ*(*t*) = *ŷ* _*p*_ (*t*) ≜ max 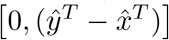 with a similar approach. For the sake of brevity, we will focus exclusively on demonstrating (7). Note that for *ξ* →0^+^, the dynamics of *ĉ* in (6) evolve on a fast timescale. In particular, let *τ* = *t/ξ* be the fast timescale, then we have 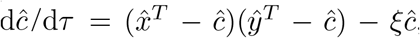, which has two equilibria that are 𝒪 (ξ) close to 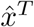 and *ŷ*^*T*^. In the following, we will prove this formally and show that *ĉ* (*t*) approaches the minimum of 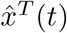 and *ŷ*^*T*^ (*t*):

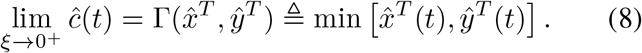

For this purpose, we define

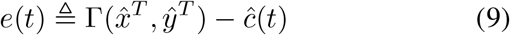

and aim to find an upper bound for |*e*(*t*)|.

**Lemma 1**. The dynamics of the error (9) satisfy *e*(*t*) ≥ 0 for all *t ≥* 0.

*Proof*. The dynamics of (3) is positively invariant in the non-negative orthant 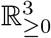. Since 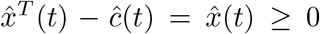 and *ŷ*^*T*^(*t*) − *ĉ* (*t*) = *ŷ* (*t*) ≥0, we have 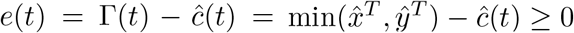.

The error dynamics can be written as:

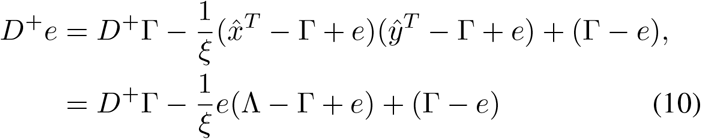

where *D*^+^ indicates right-hand derivative with respect to time and 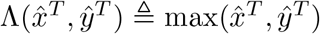. We will first show that, as a consequence of Assumption 1, the derivative *D*^+^Γ is bounded.

#### Lemma 2

Suppose that the initial conditions are such that 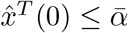 and 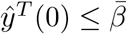, then there exist positive constants *M* and 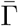, independent of *ξ*, such that 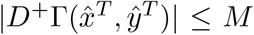 and 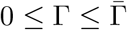 for all *t ≥* 0.

*Proof*. Consider the dynamics of the slow variables 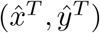 in (6) 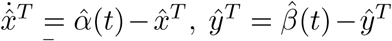. Given Assumption 1, let 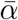 and 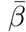 be the upper bounds for 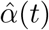 and 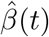, respectively. By the comparison principle, we have 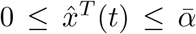 and 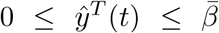 for all *t ≥* 0. Hence, we have 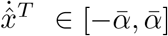 and 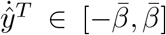. For this reason, there exists an *ξ*-independent constant *M* such that 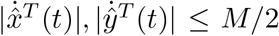 for all *t ≥*0. Because 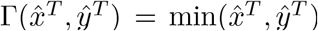, we have 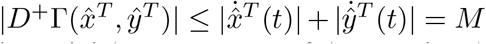. The bound on Γ(*t*) is a trivial consequence of Assumption 1.

The assumptions on initial conditions 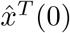 and *ŷ*^*T*^ (0) are for technical convenience and could be relaxed as the set 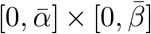 is globally attractive for 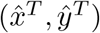. We will now use comparison principle [21] to develop bounds for the error dynamics (10):

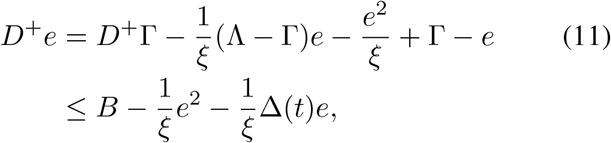

where we applied the results in Lemma 2, and 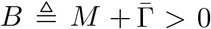 and 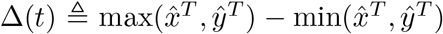 are both *ξ*-independent. The following Theorem states that, pick any *T >* 0 such that Δ(*t*) is bounded below by some *ξ*-independent constant for all *t≤ T*, there exists a sufficiently small *ξ* such that 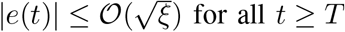 for all *t ≥ T*.

#### Theorem 1

Under Assumption 1 and suppose that there exist *ξ*-independent constants *T, δ >* 0 such that 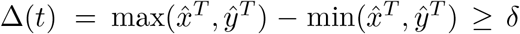 for all 0 ≤*t ≤T*. Then there exist positive constants ξ^*^ and *p*, such that

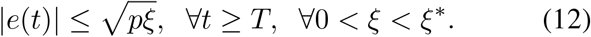

*Proof*. The proof consists of two parts. First, we show that for any *p > B* the set 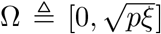 is positively invariant. This is because, by (11) and Lemma 1, for any *e ≥p*, we have *D*^+^*e ≤B −e*^2^*/ξ ≤B −p*^2^*/ξ <* 0. Hence, the set Ω is positively invariant. Next, we show that the set Ω is attractive, and for any *e*(0) *∉* Ω, there exists sufficiently small *ξ* for *e*(*t*) to enter Ω in any chosen *T*. By inequality (11), we have

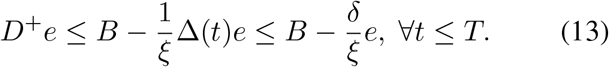

By (13) and the comparison principle [21], we have

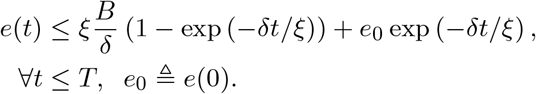

Hen ce *e*(*T*) *≤ ξB/δ* + *e*_0_ exp(*−δT/ξ*) *≤* 2*ξB/δ*, if *g*(*ξ*) ≜*ξ* ln 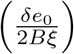 *≤ Tδ*. Since lim_*ξ→*0_+ *g*(*ξ*) = 0 and *dg/dξ*(*ξ*) *>* 0 for sufficiently small *ξ*, there exists an 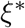 such that *g*(*ξ*) *≤ Tδ* for all 0 *< ξ ≤* 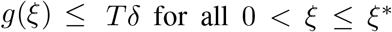. Therefore, *e(T)∈Ω*for any 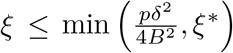. This completes the proof, as we showed that *e*(*t*) enters the positively invariant set Ω of size 𝒪 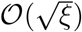 at *t* = *T* for sufficiently small *ξ*.

Because we have shown that *e*(*t*) converges to zero when *ξ →*0^+^, then expression (8) holds and so does our initial equation (7).

**Remark 1** Note that the error bound is independent of the derivative of the inputs 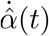 and 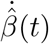. This is supported by the simulation in Fig. 2, where we varied the frequencies of the input sinusoidal signals but the error magnitude remains roughly constant. The simulation in Figure 2 shows that *𝒪* 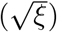 is a tight upper bound of the error dynamics. Simulations are carried out with 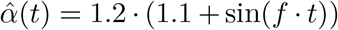 and 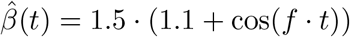.

**Fig. 2.**
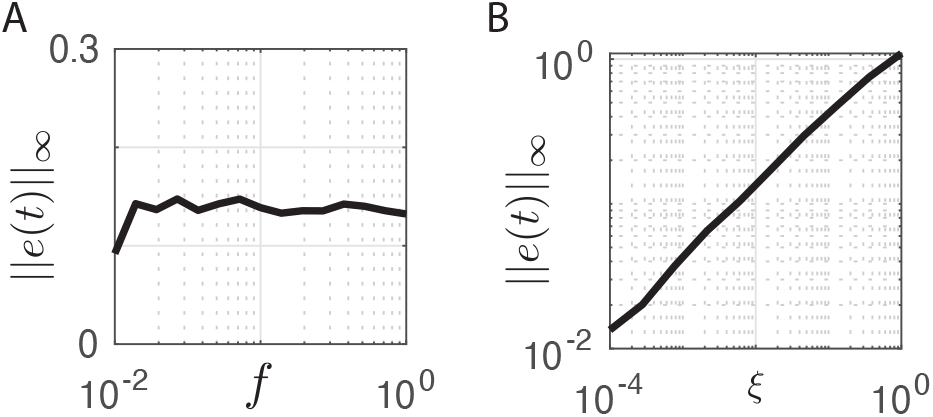
Error Analysis. (A) The magnitude of the error 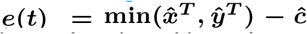 is plotted as a function of input frequency, ***f***. The simulation is carried out using the setting described in Remark 1 and ***ξ*** is kept constant at 0.01. The error is largely independent of the input frequency. (B) the error magnitude is plotted as a function of ***ξ***. The simulation is carried out using the setting described in Remark 1 and input frequency is kept constant at 0.1.

**Fig. 3.**
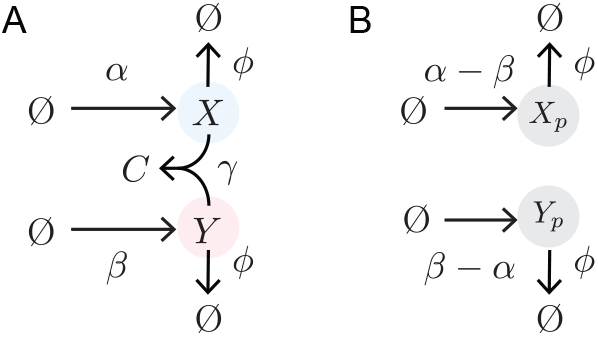
Equivalent chemical reactions. (A) The original sequestration network. (B) The equivalent reactions corresponding to the approximated dynamics (14).

**Remark 2** If the complex *C* dissociates back into reactants *X* and *Y* with reaction parameter *γ*^−^, the error analysis becomes perturbed by an additional term of order *𝒪* (*ξ*), which is negligible given the 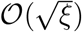 upper bound in Theorem 1.

#### Remark 3

With different degradation constants for *x, y* and *c*, equations (6) become coupled, and the analysis above cannot be easily adapted to consider this case. Computational simulations (not shown) suggest that a large discrepancy among degradation rates can deteriorate the subtraction computation.

### IV. Linear approximation and computational solution of the subtraction network

Building on the result of the previous section we can introduce a linear approximation to the dynamics of the molecular sequestration network (2). The approximation holds for *ξ →*0^+^. By taking the derivative of the right side of equation (7) (dimensional equations) and substituting the definition of 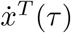 and 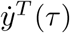, we find:

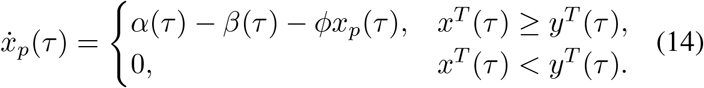

Similarly, we can derive the approximation for *ŷ*:

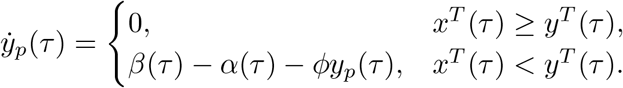

These piecewise linear ODEs include two species whose dynamics are decoupled and can be solved exactly and independently. We can also map these ODEs to an “equivalent” chemical reaction network sketched in Fig.3. When *α* and *β* are constant, the dynamics of *x*^*T*^ and *y*^*T*^ are linear (see the first two non dimensional equations in model (5)) and *x*_*p*_(*τ*) can be found exactly.

Simulations of the subtration module are shown in Fig. 4A. Specifically, the bottom panel shows that the solution *x*(*τ*) (blue shades) of the nonlinear model (2) converges to the approximation *x*_*p*_ (black) as *γ → ∞* (*ξ →* 0). When the input rates *α*(*τ*) and *β*(*τ*) are time varying, equations (14) can be easily integrated to obtain the estimates *x*_*p*_ and *y*_*p*_ without pre-computing *x*^*T*^ and *y*^*T*^ : inequality *x*^*T*^ (*τ*) *< y*^*T*^ (*τ*) can be replaced by *x*_*p*_(*τ*) *<* 0, and *x*^*T*^ (*τ*) *≥ y*^*T*^ (*τ*) can be replaced by *x*_*p*_(*τ*) *≥* 0 (similarly for *y* and *y*_*p*_).

**Fig. 4.**
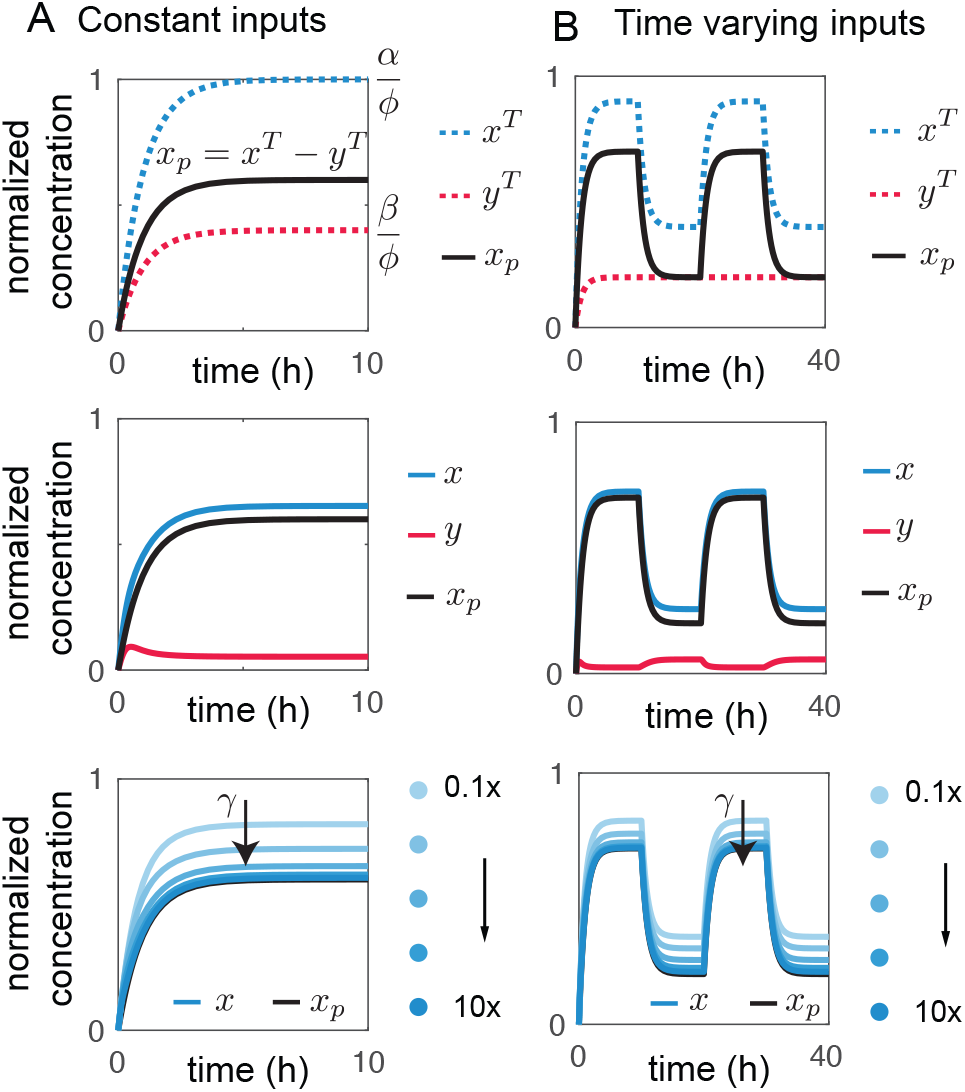
Approximated solution of molecular sequestration. Behavior of the sequestration operator under (A) a constant inputs ***α* = 1 *μM/h*** and ***β* = 0.4 *μM/h*** (***γ* = 10*/μM/h***, and ***ϕ* = 1*/h***) and (B) a time-varying square wave input where 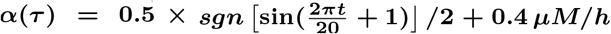 **and β(τ) = 0.2 *μ*M/h. Top** panels highlight the definition of ***x***_***p***_ **= *x***^***T***^ ***−y***^***T***^. The center panels compare the approximated dynamics of ***x***_***p***_ (black) with the simulated dynamics of ***x*** (blue) and ***y*** (red). Bottom panels simulated dynamics of ***x*** (blue) converging to the approximated dynamics ***x***_***p***_ (black) as ***γ → ∞***. These simulations integrate equations (2) and the approximation (14).

We illustrate this with the simulations in Fig. 4B. As a test of time-varying input we take *α*(*τ*) to be a square wave, while input *β*(*τ*) is kept constant. The top panel of Fig. 4B reports the trajectories for *x*^*T*^ = *x* + *c* (red dotted line), *y*^*T*^ = *y* + *c* (blue dotted line), and the approximate trajectory *x*_*p*_ = *x*^*T*^ *−y*^*T*^ (14) (black solid line). The center panel in Fig. 4B compares the trajectories of *x* (solid blue line) and *x*_*p*_ (solid black line); this plot also shows that *y* (solid red line) is small but not zero. The bottom panel of Fig. 4B compares the trajectories of *x* (solid blue lines) for different values of sequestration constant *γ* and the ideal trajectory *x*_*p*_(solid black line): larger *γ* results in the overlapping of *x* and *x*_*p*_. To summarize, for sufficiently large *γ* (which is equivalent to taking a small *ξ*) the solution computed for these linear ODEs approaches the full solution of the nonlinear ODEs.

### V. Building molecular feedback controllers through the subtraction operator

Network (2) has been used as a molecular feedback controller [22]. In this context, we re-examine the network by focusing on its capacity to perform a subtraction operation and by taking advantage of the approximations derived earlier. Our goal is to regulate a standard protein production/degradation process, described by these chemical reactions:

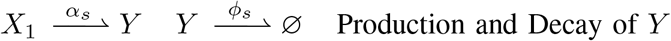

where we do not have direct control over *α*_*s*_ and *ϕ*_*s*_, while we can manipulate the abundance of species *X*_1_. We want to match the level of the process *Y* with the level of a reference species *R* (Fig. 5B and C, gray box).

**Fig. 5.**
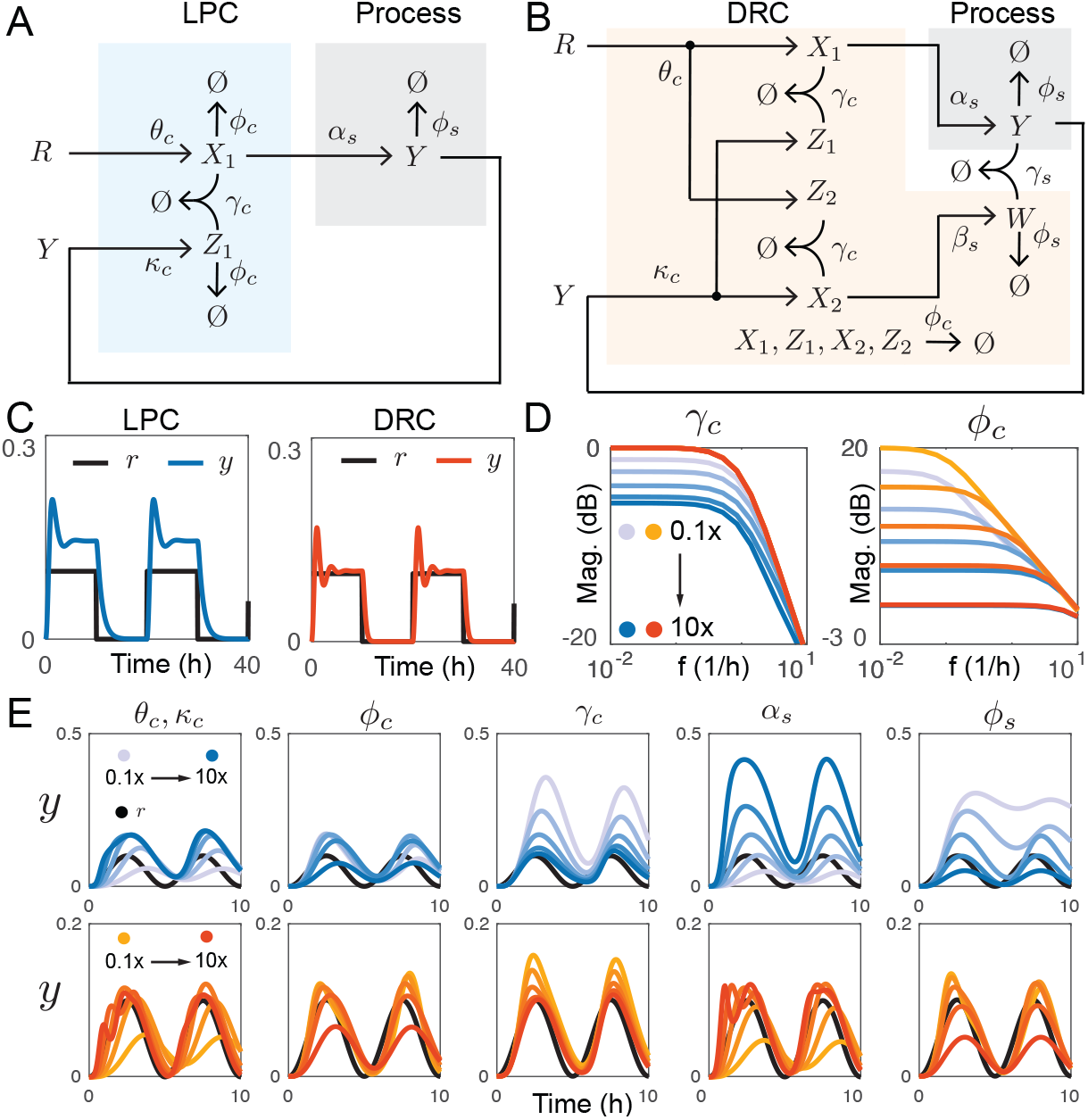
Simulations comparing controller performance. Reaction schematics of the LPC (A) and DRC (B). (C) Example trajectories under a square wave input. (D) Empirical frequency analysis. (E) Tracking a sinusoidal reference under variation of individual parameters of the controllers (***θ***_***c***_ **= *k***_***c***_, ***ϕ***_***c***_, ***γ***_***c***_) and of the plant (***α***_***s***_, ***ϕ***_***s***_). The nominal parameters are ***θ***_***c***_ **= *k***_***c***_ **= 10 */h, γ***_***c***_ **= *γ***_***s***_ **= 10 */μM/h, α***_***s***_ **= *β***_***s***_ **= *ϕ***_***s***_ **= *ϕ***_***c***_= 1 */h*

### A low-pass controller (LPC)

We begin by applying the analysis described earlier to a leaky molecular sequestration controller [5]: this exercise is useful as it brings out its inherent similarity to a low-pass controller (LPC). We introduce two controller species, *X*_1_ and *Z*_1_. The reference species *R* produces *X*_1_ at a rate parameter *θ*_*c*_; to close the loop, we use species *Y* (process output) to induce the production of species *Z*_1_ at a rate parameter *κ*_*c*_. The controller species *X*_1_ and *Z*_1_ sequester each other at a rate parameter *γ*_*c*_ (forming a complex *C*_1_), and they decay at rate parameter *ϕ*_*c*_. The reactions are summarized below (blue box in Fig. 5A):

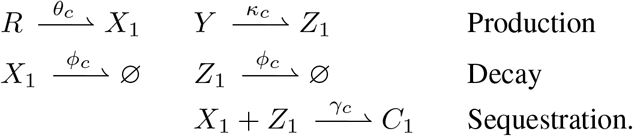

Through the law of mass action, the closed loop model is:

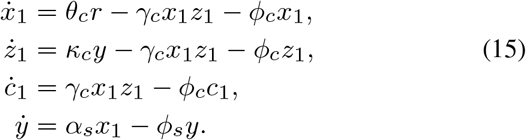

Our previous analysis (*γ → ∞*) allows us to derive a simple interpretation of how this controller works. Through a change of variable 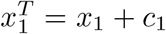, and 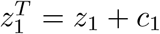 we can find the approximated solution to *x*_1_(*τ*):

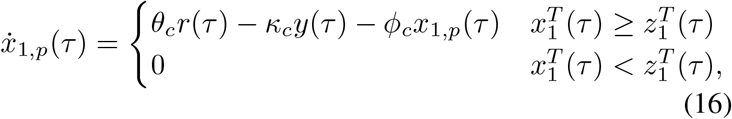

and similarly we can find the dynamics of *z*_1,*p*_. By defining the rescaled error 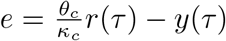, it is easy to derive the frequency response mapping *e* to *x*_1,*p*_. When 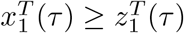:

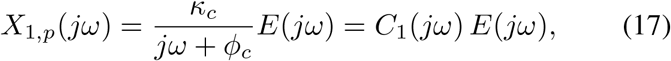

when 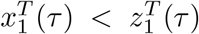, *X*_1,*p*_(*jω*) = 0. In expression (17), *C*_1_(*jω*) is an LPC with DC gain *κ*_*c*_*/ϕ*_*c*_ and corner frequency *ϕ*_*c*_. At low frequencies (*ω «ϕ*_*c*_), *C*_1_(*jω*) behaves like a proportional controller with gain *κ*_*c*_*/ϕ*_*c*_. For *ϕ*_*c*_ *→*0, i.e. in the absence of dilution/degradation, *C*_1_(*jω*) becomes an integrator *C*_1_(*jω*) = *κ*_*c*_*/jω*, consistently with dilution-free sequestration controllers [3]. It is known that ideal integral behavior is possible only in the absence of dilution/degradation [5]. In the presence of degradation, the leaky molecular sequestration controller can only achieve quasi-integral action: fast sequestration and large production rate parameters [5], [16], [18], or input-output ultrasensitivity [5], [23] are required to minimize the steady-state error between reference and output.

### A dual-rail controller (DRC)

It is apparent that the approximated output *x*_1,*p*_ of our controller (16) is non-zero only when 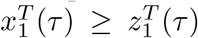, meaning that it only responds to the positive part of the difference between 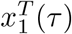 and 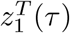. When 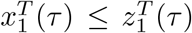, a positive response could be obtained through *z*_1,*p*_ as a second output, or by including a second controller. So we propose a “dual rail” architecture that combines two controllers in parallel and should improve the performance of the single low-pass controller. To build a dual rail controller (DRC), we connect two sequestration networks in parallel and one in series, and we show that this architecture allows a faster response when compared to a single sequestration module. This idea builds on previous work introducing dual rail molecular circuits [20] and feedback controllers [24], [25] in the context of DNA nanotechnology. The DRC is sketched in the orange box in Fig. 5B: it includes three sequestration reactions, and two outputs, species *X*_1_ that directly regulates the output *Y* (like in the single rail controller) and species *W* which removes output *Y*. The first sequestration reaction is identical to the single rail controller and produces *X*_1_. The second sequestration reaction consists of species *X*_2_ and *Z*_2_: when compare to the single rail controller, the inputs to this reaction are swapped. The reactions are:

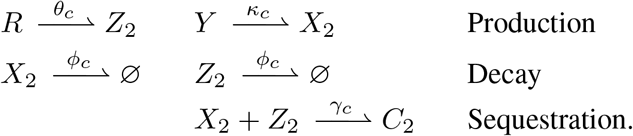

The third sequestration reaction serves as a mechanism to actively produce and remove plant species *Y*, and includes the reactions:

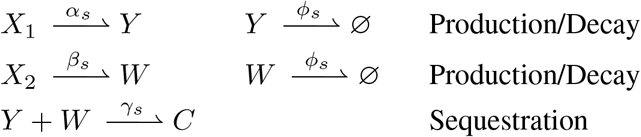

The closed loop system including the dual rail controller is:

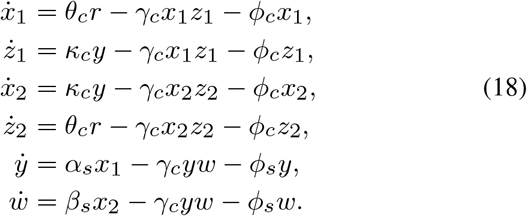

We omit the dynamics of *c*_1_ (formed by binding of *x*_1_ and *z*_1_), *c*_2_ (formed by *x*_2_ and *z*_2_) and *c*_3_ (formed by *y* and *w*).

### LPC and DRC performance comparison

Fig. 5C shows example kinetic traces of the LPC and DRC, showing that the LPC tracks the reference with a larger steady state error. This error may due to the presence of dilution *ϕ*_*c*_ ≠ 0 (non-ideal integral controller), or to poor error computation -intended here as the computation of the positive part of the error only. This controller can actively increase *y* (positive action), while only dilution/degradation can decrease the amount of *y*. This type of control strategy with just positive action leads to a trade-off between the controller dynamics (that increases *y*) and the process dynamics (that removes *y*). When *ϕ*_*c*_ ≠ 0 the gain *κ*_*c*_*/ϕ*_*c*_ decreases, contributing to a larger steady state error. Even in the presence of dilution, the DRC shows a reduced error presumably because it actively increases and reduces *y* (using the full error computation).

Fig. 5D shows the “empirical” frequency response of the two controllers as the sequestration rate *γ*_*c*_ or the decay rate *ϕ*_*c*_ are varied. The controller equations in the LPC model (15) and DRC model (18) were fed a periodic input reference *θ*_*c*_*r* = 0.5(sin(*fτ*) + 1)*/*2 + 0.3 *μM/h*, and fixed *κ*_*c*_*y* = 0.3 *μM/h* (we compute the frequency response of the controller in isolation, assuming a constant level of *y*). We are not able to provide an exact form of the DC gain for the DRC, due to the complexity of its equations.

We then numerically integrated the equations, and measured amplitude and frequency of the output *x*_1_ for the LPC, and the difference of *x*_1_ −*x*_2_ (these species deliver opposite actuation on the plant) for the DRC. In these simulations (blue for the LPC, orange for the DRC) we vary either the sequestration rate *γ*_*c*_ or the decay rate *ϕ*_*c*_ parameter. Both plots resemble the Bode plot of an LPC with a corner frequency near *ϕ*_*c*_. In the left panel (*ϕ*_*c*_=1, *γ*_*c*_ varies), the “DC gain” of the DRC does not depend on the sequestration constant *γ*_*c*_ as the LPC. In the right panel, (*ϕ*_*c*_ varies, *γ*_*c*_ = 10) the corner frequency moves as the value of *ϕ*_*c*_. There is a small offset between the LPC and the DRC, that is inversely proportional to *ϕ*_*c*_: as *ϕ*_*c*_ decreases, passive down-regulation is reduced and the DRC performs better than the LPC.

Finally, Fig. 5E illustrates the closed loop performance when the reference is a sinusoid, as individual parameters are changed. We assume *θ*_*c*_ = *κ*_*c*_ in all cases. Larger values of *θ*_*c*_ = *κ*_*c*_ improve the tracking performance of both LPC and DRC, however the LPC always shows a significant larger error. Increasing or decreasing *ϕ*_*c*_ in the LPC does not improve its tracking behavior, while for *ϕ*_*c*_ *→*0 the DRC shows error reduction. When *γ*_*c*_ is varied, even at small values the DRC maintains better tracking when compared to the LPC. When the plant parameter *α*_*s*_ is perturbed, the DRC confirms to be better at rejecting this disturbance. When changing the plant dilution constant *ϕ*_*s*_, the DRC performance improves for smaller values, while the LPC performance deteriorates.

## VI. Conclusion and discussion

We have described an approximation of the dynamic behavior of molecular sequestration that illustrates how sequestration computes the difference between input signals.

When compared to previous theoretical work on this topic, our analysis includes dilution/degradation of the reactants (which is always present in living cells) and considers time-varying inputs. The approximation we derived is useful to illustrate how leaky molecular sequestration controllers can be considered low-pass controllers that perform only one side of the subtraction operation. Through simulations, we showed that a dual rail controller (DRC) performs a full subtraction operation, and achieves a lower tracking error across a range of parameters. The DRC requires three sequestration reactions, one of which involves the output *Y* and species *W*. In practice, this may be realized through *de novo* protein switches [26], or *σ* factors (*Y*) binding with anti-*σ* factors (*W*) driving *σ*-responsive promoters to express *X*_2_ and *Z*_1_ [18]. The reactants *X*_*i*_ and *Z*_*i*_ (*i* = 1, 2) may be realized introducing RNA-RNA interactions [16], [17]. We expect that our study will accelerate and generalize the design of biomolecular controllers.

## Notes

This work has been partially supported by the BBRSC-NSF/BIO award 2020039 and by NSF CCF/SHF award 2107483 to EF

https://ieeexplore.ieee.org/document/10180067/authors#authors

### Competing Interest Statement

The authors have declared no competing interest.

### Summary of Updates

We simplified the model to make the derivations simpler.

